# Inter-Individual Variability in Dorsal Stream Dynamics During Word Production

**DOI:** 10.1101/2022.04.01.486723

**Authors:** C. Liégeois-Chauvel, A.-S. Dubarry, I. Wang, P. Chauvel, J.A. Gonzalez-Martinez, F.-X. Alario

## Abstract

The current standard model of language production involves a sensorimotor dorsal stream connecting areas in the temporo-parietal junction with those in the inferior frontal gyrus and lateral premotor cortex. These regions have been linked to various aspects of word production such as phonological processing or articulatory programming, primarily through neuropsychological and functional imaging group studies. Most if not all the theoretical descriptions of this model imply that the same network should be identifiable across individual speakers. We tested this hypothesis by quantifying the variability of activation observed across individuals within each dorsal stream anatomical region. This estimate was based on electrical activity recorded directly from the cerebral cortex with millisecond accuracy in awake epileptic patients clinically implanted with intracerebral depth electrodes for pre-surgical diagnosis. Each region’s activity was quantified using two different metrics—intra-cerebral evoked related potentials and high gamma activity—at the level of the group, the individual, and the recording contact. Using picture naming task, the two metrics show simultaneous activation of parietal and frontal regions in line with models that posit interactive processing during word retrieval. They also reveal different levels of variability across brain regions and patients except in auditory and motor regions. The independence and non-uniformity of cortical activity according to the two metrics push the current model towards sub-second and sub-region explorations focused on individualized language speech production. Several hypotheses are considered for this within-region heterogeneity.

## Introduction

In human cognitive neuroscience, linking a processing model with single-case patient data provides a fruitful means to pursue improvements of both the model and the clinical assessment. Constructing “typical” participants from aggregated (e.g. average co-registered) data has been the primary focus of research on brain function (Seghier & Price, 2018), but there is increasing interest in understanding various kinds of inter-individual differences. Stable and interpretable individual variability in resting state neural networks has been repeatedly observed, pointing out the existence of spatially variable “network variants” (Gordon et al., 2017; D’Esposito 2019; Seitzman et al. 2019). The potential sources of variability in brain function during task performance have been conceptualized and evidenced across various cognitive domains (Miller et al., 2009; Sanfratello et al., 2014; for a review, see Seghier & Price, 2018)

The neural network involved in language processing is broad and known to be variable across healthy individuals (Mahowald & Fedorenko, 2016) and in neurological disorders (e.g. Berl et al., 2014). This inter-individual variability has been frequently studied to understand variations in the hemispheric specialization for language (Tzourio-Mazoyer et al., 2017). It has also been harnessed to distinguish a “core” part of the language network from regions recruited across other cognitive domains (Fedorenko et al. 2011; 2012), or to explore how a given task (e.g. word reading) might be achieved through more than one processing pathway (Seghier et al., 2008). This research has been mostly concerned with perceptual processes (e.g., recognizing words and their meanings) or with higher levels of abstraction (e.g., comprehending sentences). In contrast, language production theories in cognitive neuroscience (Hickok, 2012; e.g. Indefrey, 2011; Roelofs, 2014; Strijkers & Costa, 2016) have been mostly developed and tested under the (often implicit) hypothesis that a roughly similar network, with roughly similar spatial or temporal characteristics, must be present in every individual. Consequently, the interactions between language production models and individual patient data remain cautious. One representative example is the exploration of the language production network in the context of epilepsy surgery (Arya et al., 2018; Benjamin et al., 2020; Nakai et al., 2019; Trebuchon et al., 2020).

According to current models, producing a word to express a particular thought requires that a single item is selected among alternatives, and that its form is encoded to be articulated overtly (Levelt et al., 1999). These cognitive processes are sub-served by diverse functionally specialized regions (Indefrey, 2011). Their network intricacy has been clarified in terms of the dual stream framework for language processing with a distinction between (i) the ventral stream, a temporal-lobe stream which maps semantic knowledge on to lexical representations (e.g. during visual naming), and (ii) the dorsal stream, a parieto-frontal stream which interfaces auditory-phonological information with the motor system (Hickok, 2012; Roelofs, 2014; Ueno et al., 2011).

The dorsal stream we focus on is thought to be left lateralized in most right-handed individuals (Gleichgerrcht et al., 2015; McKinnon et al., 2018; Saur et al., 2008; Schwartz et al., 2012; Warren et al., 2005)(but see bilateral processes reported in Cogan et al., 2014). Between the temporo-parietal and lateral frontal cortices, the dorsal stream is supported by two of the anatomical pathways that form the superior longitudinal fasciculus (SLF). Its deep segment connects the posterior temporo-parietal regions to the Pars Opercularis while the lateral part of SLF (SLF III) links the Supra Marginal Gyrus (SMG) to the ventral premotor cortex (Catani & Mesulam, 2008; Parlatini et al., 2017; Petrides, 2014). Other regions that form the dorsal stream, such as Supplementary Motor Area (SMA) and Ventral Motor Cortex, are also connected to posterior parietal areas via the dorsal branch of the superior longitudinal fasciculus (SLF I).

This network is swiftly activated during word production, with responses typically occurring within a second in experimental settings such as picture naming. The corresponding cognitive operations and neural activations are thus coordinated on a sub-second scale. An influential theoretical proposal combined a cognitive model with a meta-analysis of imaging and electrophysiological data into a framework that estimates a cascade of onsets for the different regional activations and processes (Indefrey, 2011). An alternative proposal hinges on hierarchical processing and top-down modulations, thereby highlighting the sustained reverberatory activity during which diverse processes largely overlap in time (Rapp & Goldrick, 2000; Strijkers & Costa, 2016).

An assessment of inter-individual variability in dorsal stream activation must thus be conducted with sub-second temporal resolution. Magneto-encephalographic (MEG) signals have a fair spatial resolution and a high temporal resolution, but synthetizing them in a unified spatio-temporal model of word production has proved to be challenging (Munding et al., 2016). The cortical dynamics underlying word production are increasingly explored in studies with invasive recordings (e.g. Nakai et al., 2019; Piai et al., 2016; S. K. Riès et al., 2017)(for review, see Flinker et al., 2018; Llorens et al., 2011). But these studies typically acknowledge the variability across patients without seeking to quantify or interpret it beyond its use for computing statistics at the group level (Flinker et al., 2018; Lachaux et al., 2012). The hypothesis is that such central tendency measures will erase possible pathological idiosyncrasies (Lachaux et al., 2012; Parvizi & Kastner, 2018), with the associated risk of losing important information or distorting data patterns (Seghier & Price, 2018).

## The current study

Despite increasing theoretical interest for inter-individual variability in neural functional activity, and its importance for translating between processing theories and beside assessments of the language network, inter-individual variability in the language production network remains under-studied. Here, we explored the variability in the activation of fronto-temporo-parietal regions that encompass the dorsal stream (Hickok, 2012), when they were recruited for word production, a task that is typically described to be solved with a single processing pathway (Indefrey, 2011). At stake in our approach was whether the variability described for networks in other contexts (D’Esposito, 2019; Gordon et al., 2017; Seitzman et al., 2019) could be identified at the level of the individual regions implicated.

The quantification of inter-individual variability was performed on a dataset obtained from epilepsy patients involved in pre-surgical diagnosis procedures (StereoElectroEncephalography or SEEG) that required the implantation of intra-cerebral depth electrodes. Recordings from these electrodes gave us access to the variability in time-resolved electrophysiological signals rather than the spatial variability derived from functional magnetic resonance imaging (fMRI) data (Berl et al., 2014). A link between signals in the high gamma frequency band and cognitive processes has been demonstrated in a variety of experimental tasks (Lachaux et al. 2012). However, high gamma activity (HGA) does not capture all of the neurophysiological modulations (Flinker et al. 2018), as shown in intra-cerebral recordings during auditory perception (Edwards et al., 2009; Nourski et al., 2015), or in MEG recordings during our task of interest, picture naming (Laaksonen et al. 2012). Thus, we decided to investigate both HGA modulations and intra-cerebral evoked related potentials (iERP). We explored variability across patients but also across single-electrode time courses, looking to assess the diversity of activities within each region of interest.

## Methods

### Patients

Epileptic patients were recruited from the pool of patients assessed by the Cleveland Clinic Epilepsy Center for surgical treatment of medically refractory Epilepsy. They were undergoing a stereo-electro-encephalography (SEEG) diagnostic evaluation as part of their pre-surgical assessment. (Bancaud et al., 1969; Chauvel et al., 2019). After an anatomo-functional localizing epileptogenic network assumption is formulated, a strategy to implant depth electrodes to specific regions of the brain is defined. Depth electrodes are stereotactically inserted using a robotic surgical implantation platform (ROSA, Zimmer-Biomed, USA), in a three-dimensional arrangement with either orthogonal or oblique orientations, allowing intracranial recordings from lateral, intermediate or deep cortical and subcortical structures (González-Martínez 2015). For each patient, between 8 and 13 stereotactically placed electrodes were implanted. Within each electrode, the recording contacts were 2 mm long, 0.8 mm in diameter, and spaced 1.5 mm apart. The insertion of these electrodes is based purely on clinical needs and is made independently of any research related purpose. Part of the procedures seeks to identify functional networks such as those involved in language, to be able to spare them from the surgical procedure (Trebuchon et al., 2020).

We studied 17 epileptic patients who were native speakers of English. Among them, 11 were implanted unilaterally (9 in the left and 2 in the right hemispheres, respectively), and 6 were implanted bilaterally (i.e., they had electrodes implanted in both hemispheres, but not necessarily in directly homologous areas). Patients were invited to participate in a picture naming experiment when their implantation included depth electrodes within the parieto-frontal networks known as the dorsal language stream (Hickok, 2012). All patients enrolled voluntarily after giving written informed consent under criteria approved by the Cleveland Clinic Institutional Review Board (N°13-1248). Patient details are listed in Table 1. One patient (P15) was unable to complete the cognitive task properly, thus the analysis involves data from only 16 patients.

### Anatomical reconstruction of electrode positions

#### MRI acquisition

All MRI scans were acquired from a 3T Siemens Skyra scanner (Siemens, Erlangen, Germany). The T1-weighed Magnetization Prepared Rapid Acquisition with Gradient Echo (MPRAGE) volumetric scan was used for co-registration with CT. *MRI and CT fusion.* Immediately after SEEG implantation, a high-resolution stereo-computerized tomography (CT) was taken to obtain the anatomical location of the implanted electrodes. The post-implantation CT scan was exported into Curry 7 (Compumedics Neuroscan, Hamburg, Germany), and co-registration with the MPRAGE images was performed using an automated full-volume registration with maximization of mutual information. The accuracy of the co-registration was inspected visually and confirmed in all patients by a neurosurgeon (JGM) to ensure accuracy. The location of each electrode contact, determined by the center of the highest intensity on the CT, was individually labeled and superimposed on the MRI for visualization of its anatomical location.

#### Normalization of MRI and CT

Further processing was performed within SPM12 (Wellcome Department of Cognitive Neurology, London, UK) in MATLAB 2015a (MathWorks, Natick, Massachusetts) to normalize the locations of the electrodes of interest from all the patients. Three processing steps were performed: (1) co-registration of the CT to the MRI for each individual patient; (2) normalization of the individual MRI with the Montreal Neurological Institute (MNI) 3D common stereotactic space (Collins et al., 1998) (3) normalization of the co-registered CT in step 1 to the MNI space by applying the same transformation matrix as obtained in step 2. All the normalized CT and MRI images were then fused in Curry 7, so that the electrodes could all be superimposed on the normalized MRI and visualized on a template according to the Talairach stereotactic coordinate system.

Given the focus of this research, our exploration was limited to regions located within the language dorsal stream (see Introduction) and organized in seven anatomical groups (Figure 1). Each contact (N = 447) was localized to a specific brain region of the Talairach atlas (Talairach & Tournoux, 1988) based on its coordinates and confirmed by individual visual verification. Only regions which were sampled in at least 3 patients were included in the analysis, to allow assessing cross-patient consistency of neural responses, as described below.

**Figure 1.**
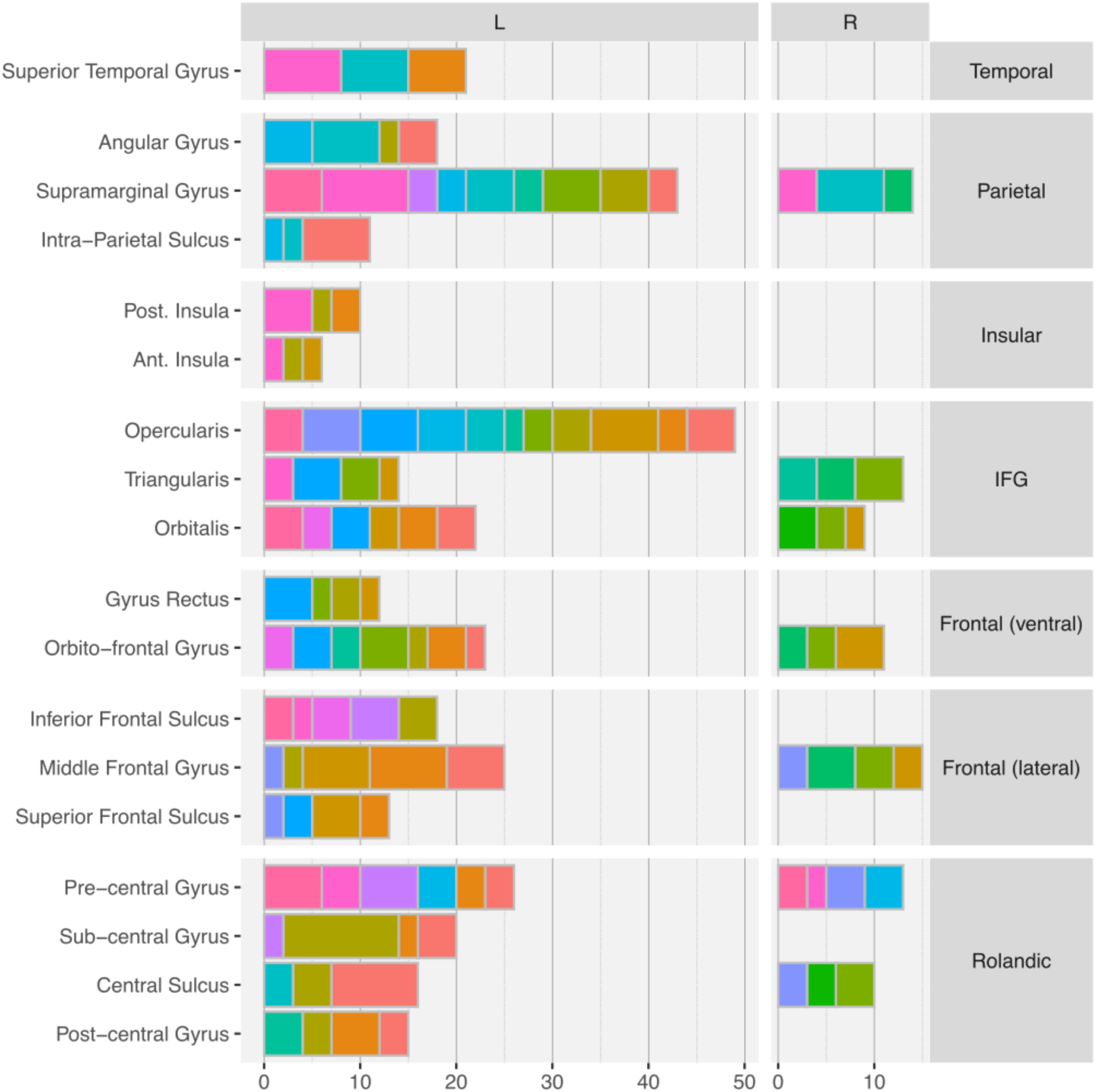
Anatomical sampling for 16 patients. Each horizontal bar represents the number of recording contacts that were identified to lie in each of the brain regions of interest. Each color is a patient. For example, Superior Temporal Gyrus was sampled 21 times across three patients, each contributing 8, 7, 6 contacts, respectively.

### Experimental task

Patients were asked to name out loud pictures of common objects from the Snodgrass and Vanderwart (1980) collection. There were 36 common items, chosen from six different semantic categories (Accessories, Buildings, Kitchen Utensils, Fruits, Furniture, and Musical Instruments). The pictures were presented in short blocks each involving six items randomly repeated five times, yielding 30 trials per block. A total of 8 blocks was planned for each patient (i.e., 240 trials), but not all patients completed all the blocks due to clinical circumstances (e.g., fatigue or interruptions). The items within a block could either be from a single semantic category (semantically homogeneous blocks) or from six different semantic categories (semantically heterogeneous blocks). This design and materials were reused from a previous research conducted with French speakers (Anders et al., 2019; Llorens et al., 2016), but with different patients completing different numbers of blocks, the homogenous vs. heterogeneous contrast was unbalanced and could not be explored in the current analysis.

The experiment was controlled by the software E-Prime v2.0.1 (Psychology Software Tools, Pittsburgh, USA). The pictures were presented on the center of the screen within a visual angle of 6° × 6°. For a subset of the patients (N=8), naming latencies were recorded with a microphone (audio Technica ATR20) placed about 13 cm in front of the patient. In this case, response times were automatically recorded in milliseconds by the software’s voice key. A trial consisted of a fixation point (variable duration across trials, between 1400 and 2100 ms), followed by the black and white target picture (presented for fixed duration of 1000 ms). Various pseudo-random orders were created to vary across patients the order of items within a block and the order of the blocks within the experiment.

### Data acquisition

SEEG electrophysiological data was acquired using a video-EEG monitoring data acquisition system (Nihon Kohden 1200, Nihon Kohden America, USA) at a sampling rate of 1000 Hz per channel. During the cognitive task, stimuli and behavioral event data were simultaneously acquired along with the SEEG signal (Johnson et al., 2014), and stored for subsequent analysis. The recordings were on-line referenced to a scalp-electrode located on vertex (Cz).

### Data analysis

Off-line pre-processing was performed in Brainstorm (Tadel et al., 2011), which is freely available for download online under the GNU public license (http://neuroimage.usc.edu/brainstorm). Evoked potentials and HGA were computed and combined into a group analysis using the Multi-patient Intracerebral data Analysis toolbox (MIA: Dubarry et al., 2022).

For each participant, the continuous monopolar SEEG recording was filtered digitally (FIR high pass filter implemented in Brainstorm; lower cutoff frequency: 0.5 Hz; stop band attenuation: 60dB). Epochs starting 1500 ms before stimulus onset and lasting 1500 ms post-stimulus were created, for a total of 3000 ms per trial. Only the epochs corresponding to correct behavioral responses were kept for the analysis. Epochs containing interictal epileptic activity (spikes or abnormal sharp waves) were removed manually following visual inspection of all epochs.

#### Intra-cerebral evoked related potentials (iERP)

The signals recorded with the Cz electrode as reference (monopolar montage) were averaged across trials to yield iERP. iERP arise from synaptic activity of large numbers of neurons distributed over a cortical region. They primarily capture low frequency components and are time and phase-locked to the stimulus. They provide a measure of the synaptic input to, and the local processing within, the area where they are recorded (Lopes da Silva, 1991). Because the signals are time-locked to the stimuli, the recordings provide information about the temporal components of the underlying generators. Any local field potential can be recorded either as positive or negative response, and this polarity depends, among other things, on the location of the electrode contact with respect to the generator. This property of iERP can be conceived of as a limitation because it complicates averaging procedures across contacts and patients (see below *Within-region variability across contacts and patients* for the adopted strategy). In addition, they do not distinguish excitatory from inhibitory currents (Buzsáki et al., 2012; Lopes da Silva, 1991).

#### High gamma activity (HGA)

The second step of data analysis focused on high-gamma band (HGA) which is characterized as broadband activity measured here between 80 and 120 Hz. To estimate HGA, a bipolar montage was computed in which the activity recorded at adjacent contacts was subtracted to yield bipolar channels (e.g., B2-B1, B3-B2, etc.) Morlet Wavelet transforms were first performed separately at five consecutive frequencies between 80 and 120 Hz with a 10Hz step (i.e., 80, 90, 100, 110, 120 Hz). Then, the five resulting time courses were separately normalized with a z-score against the baseline (−800ms to −50ms) and finally summed. This procedure was used to compensate for the larger contribution of low frequencies that would otherwise overpower the contribution of higher frequencies, due to the 1/f distribution of signal power (Bédard & Destexhe, 2009) HGA has been linked to BOLD signal of fMRI (Logothetis et al., 2001) as well as neuronal firing rate (Ray & Maunsell, 2011)localized in restricted cortical areas (Lachaux et al., 2012; Mukamel & Fried, 2012). One limitation of HGA is its power, between 100 and 1000 times lower than that of the iERP (Lachaux et al., 2012), which can make it more difficult to detect. Also, partly due to the time-frequency transformation process, it is less sensitive to small variations in the timing of behavioral and neural responses than iERP are. Oscillatory burst may suffer from latency jitters, turning the relationship with the stimulus onset fairly loose (Tallon-Baudry & Bertrand, 1999).

#### Task related significant activity

Task-evoked responses were determined to be significant using a t-test across trials at each time sample. To address the resulting multiple-comparison problem in the time domain, a minimum duration threshold *T* was estimated with a bootstrap procedure. Each step of this procedure involved the random selection of the same number of trials as in the original data set, with resampling, and the identification of periods with significant activity within the baseline window (−800 ms to −50 ms) in that sample. At each iteration, the significant period of maximal duration (*p <* 0.001) was pooled into a bootstrap surrogate distribution constructed through 1000 iterations of the procedure. For each patient, the significance threshold (minimum duration of consecutive significant *t*-values, *p* < .001) was defined as the right-tail 95% quantile of that bootstrap distribution of durations. This yielded a threshold of significance for each channel (p<0.001; corrected for multiple comparisons in time) which was used to determine task related significant activity. Independently of the statistical analysis, an expert neurophysiologist (CLC) examined the responses for each contact and identified no discrepancies between the detections based on human expertise and the statistical method.

The analysis described above was conducted per patient. We also report the time courses of each patient in each region, obtained by averaging together all the contacts of a patient within a region that showed significant activity (any activity on the mask of significance). Finally, a group analysis was performed by computing the grand average of patients’ time-courses per region.

#### Within-region variability across contacts and patients

The variability of neural activities within each region of interest was estimated with a Pearson correlation metric computed from the recorded signal time-courses. Specifically, we computed the pair-wise correlations across all signals within one region, at two different levels: contacts/channels and patients. The signal was first averaged across trials to yield a time course for each contact/channel. These time courses were correlated pairwise, yielding a correlation/covariance matrix whose values were averaged. The resulting value is the “correlation across contacts” in the region. Then, the contacts of each patient were grouped, and their time courses averaged to yield a regional time course for each patient. The correlation matrix of these time courses was computed and averaged, as before, to yield the “correlation across patients” in the region.

Importantly, for LFPs, any aggregation of signals from various contacts into a single metric must consider the arbitrary sign of the evoked potentials. Typically, two contacts recording the same dipolar source activity from opposite sides would show signals of opposite polarity, an inversion that can occur within or across patients. Conversely, opposite polarity alone does not warrant a single underlying source, as the two signals may arise from different regional sources. Here, for simplicity, we did not attempt to solve the SEEG inverse problem, which might have clarified this issue, but which is very rarely explored in SEEG (Caune et al., 2014). The iERP matrix of correlation values was thresholded separately for positive and negative values, based on a heuristically determined threshold (here, | thr | = 0.7). We counted the number of high negative correlations (i.e., below -thr) and picked the channel with the highest number of anti-correlated channels. All channels highly correlated with this extreme channel, itself included, were sign flipped and the correlation matrix recomputed. Such sign flipping procedure was preferred over working on absolute valued time-courses, to preserve the shape of the signal and the information carried by potential inversions. The same analysis was conducted with other threshold values (0.6, 0.8, and 0.9) for comparison. With threshold 0.7, the number of regions that showed significantly correlated iERP was 11 out of 25 (see Results). For the other values, it was 7, 8, or 9 regions out of 25. In other words, our choice of threshold is the most liberal one. It might err on the side of showing within–region correlations, where we wanted to highlight the opposite.

To note, the sign flipping procedure was applied before patient and region averaging described above under *Intra-cerebral event-related potentials*, and only for iERP signals. In HGA, the sign meaningfully indicates activation or possibly deactivation with respect to baseline, and hence is informative. Therefore, for HGA, the averaging and the variability metric were computed without applying the sign flipping procedure.

The statistical significance of correlations by contacts/channels was assessed with a permutation test that preserved the participants’ relative contribution to each region. For any given region, we preserved the patients’ identities and number of contacts (or channels in HGA) when permutating each signal with a signal from any other brain region. The two correlation indexes described above were computed. After 1000 permutations, we obtained surrogate distributions of correlation values (two for each region, for iERP and for HGA) with which significance was established at an alpha level of 0.05.

For any region, we preserved the number of patient and number of contacts and the statistics consisted in randomly permutating these signals with those of other anatomical regions.

## Results

### Overview

All patients except P-15 completed the task correctly. This left the data from 16 patients for the analysis. Because of technical issues, response times (RT) could only be recorded in 8 out of 16 patients. In addition, the experimental software we used (E-Prime 2) detected vocal response onsets automatically based on sound intensity, without generating a recording of the sound wave for off-line checking. Such intensity based automatic detections are considered suboptimal (https://doi.org/10.3758/s13428-017-1002-7). For these reasons, we did not perform any detailed analyses based on response times; in particular, we did not use these estimates to compute response-locked activities. The average RT was 894ms (sd 105ms). Mean accuracy was above 90 %. Only trials with correct responses were included in the neural signal analysis.

The anatomical sampling criteria lead to the inclusion of 25 different regions, 18 of which were in the left hemisphere and 7 in the right hemisphere. Across patients, there were 88 different electrode probes, within which 362 electrode contacts were in the left hemisphere, and 85 in the right hemisphere (Figure 1).

### Intra-cerebral responses

The analysis of intra-cerebral evoked response potentials (iERP) for the 447 monopolar contacts revealed 359 contacts for which there was significant post-stimulus task-related activity (i.e., 80%), and 88 contacts without significant response. There was a 100% consistency between the active / inactive classification performed visually (and independently) by the expert neurophysiologist (CLC) and by the automated statistical method. For the High Gamma Activity (HGA), the 447 monopolar contacts were re-coded as 367 bipolar channels. Among them, 211 revealed significant post-stimulus task-related HGA (i.e., 58%), and 156 contacts were without significant response.

Across the 25 sampled regions, there was a strong similarity between the variability of neural responses estimated in terms of correlations between contact averages vs. correlations between patient averages (Figure 2A). This suggests high within-patient correlations. Conversely, there were substantial variations between regions and, furthermore, between correlation estimates based on iERP or HGA time courses. There were 11 regions that showed significantly correlated iERP, also 11 regions that showed significantly correlated HGA, but one significance did not necessarily imply the other (Figure 2B).

**Figure 2.**
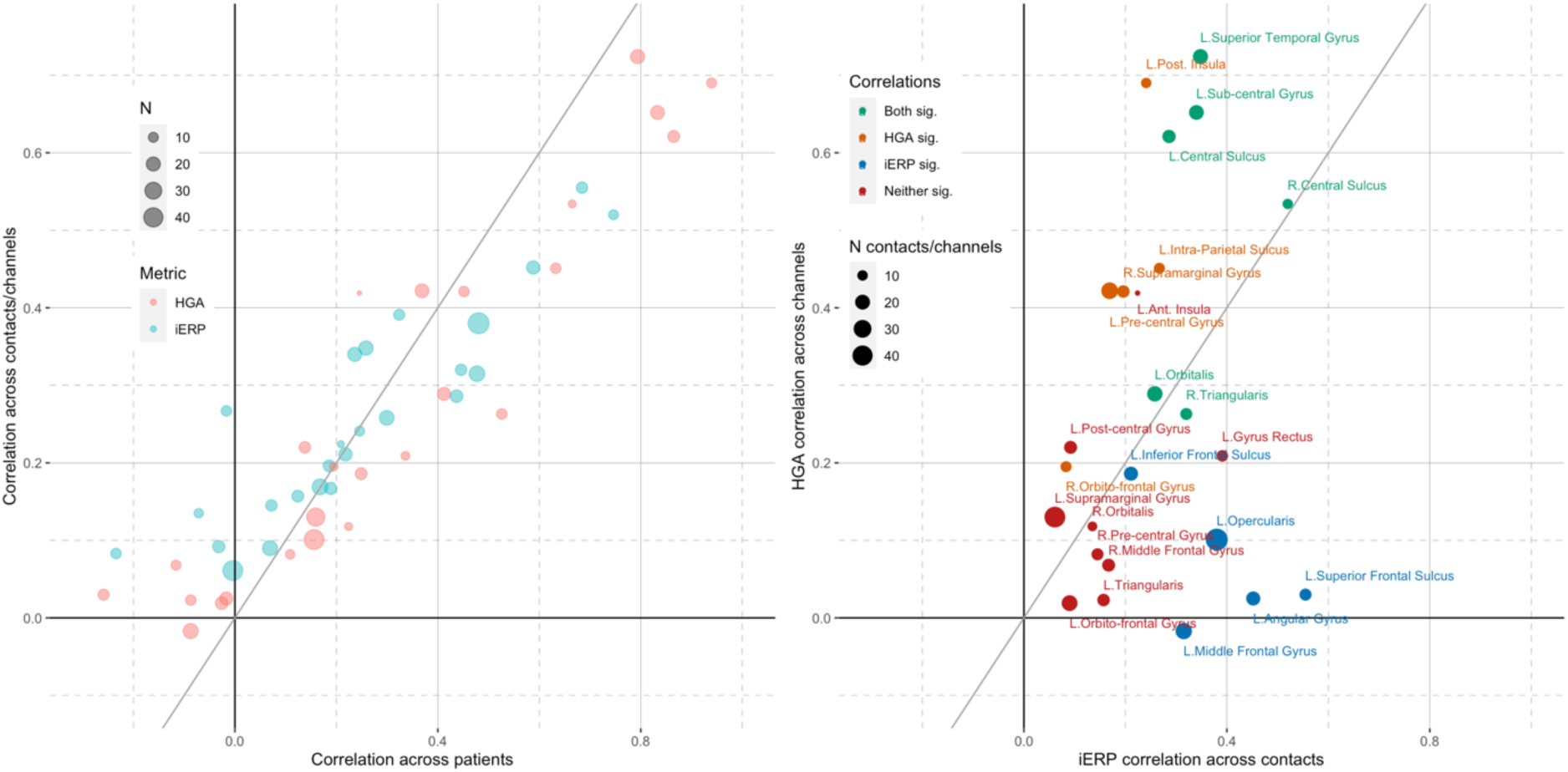
Correlation of neural responses across the anatomical regions of interest. **A.** Scatterplot of within region correlations computed on the time-courses averaged per patient (x-axis) vs. averaged per contact (y-axis), based on iERPs (red) and HGA (green) for the different ROIs (dots). Each dot is an ROI. The metrics computed across patient and across contacts yield very similar results, for both iERP and HGA. **B.** Focus on correlations across contacts, now directly compared for iERP (x-axis) vs. HGA (y-axis) time courses.. Each dot is an ROI. The two metrics show substantial disparities across regions: Both sig = both metrics show a significant correlation across contacts; HGA sig. = only HGA does; iERP sig. = only iERP does; Neither sig. = neither metric shows a significant correlation across contacts.

Regions with high correlations (i.e., small variability) are easily understood as being recruited by the task, and the consistently recorded activity can be described as the signature of the cortical region for this task. For regions exhibiting higher variability of responses, more detailed interpretations (e.g., sub-regional distinctions) will be considered. Four groups of regions were distinguished based on the significance level of variability observed with each of the two metrics (color coded in Figure 2B). Figure 3 illustrates one representative region of each of these groups.

**Figure 3:**
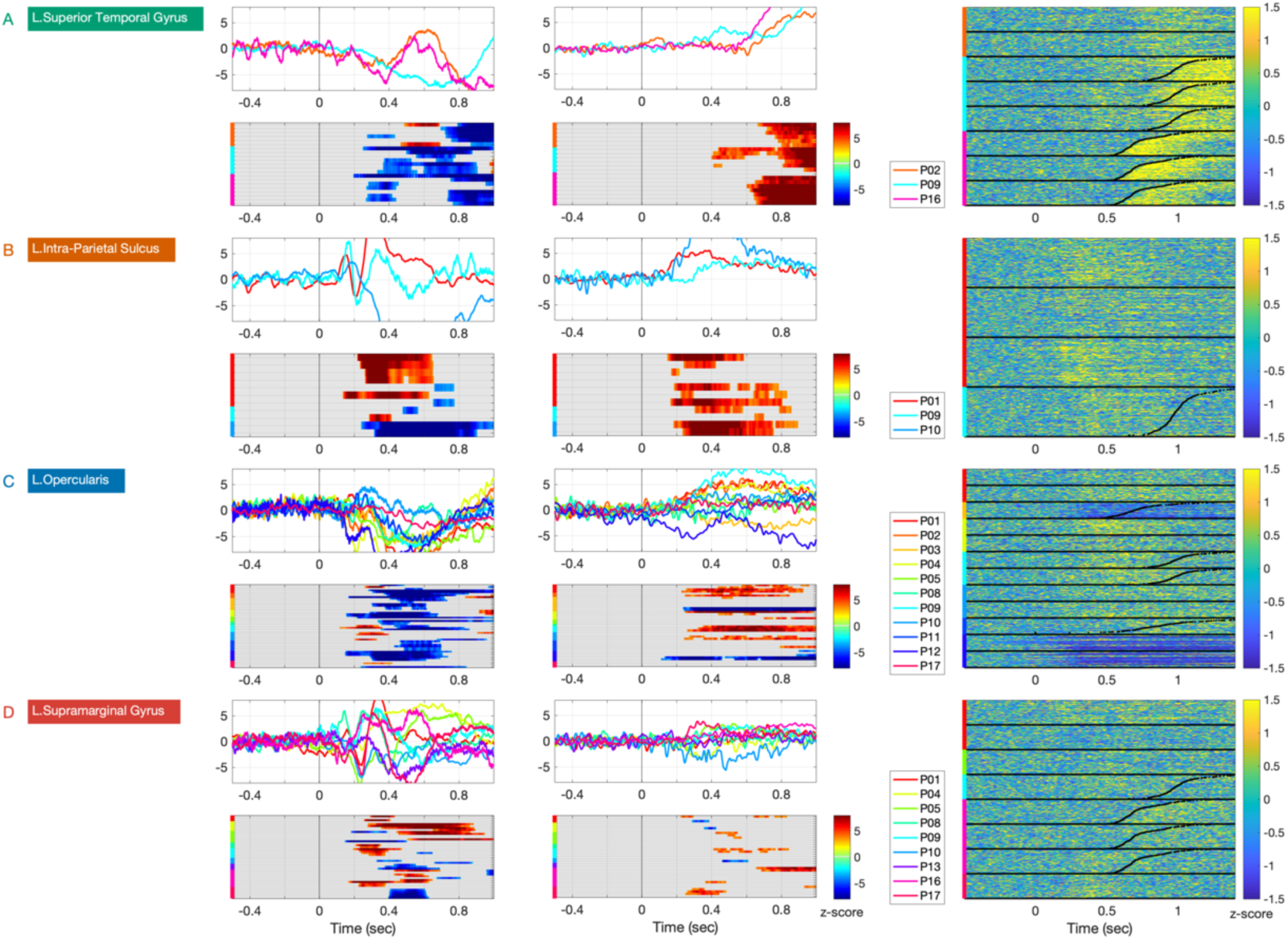
Different levels of inter-patient variability of intracerebral activity during word production in four representative regions of the left hemisphere. Left column: iERP time courses and their significance mask; middle column: HGA time courses and their significance mask in z-score against baseline; right column: single trial activity in HGA in z-score against baseline, black dots represent response times (where available). The color of the region names boxes corresponds to those in Figure 2b. A-Superior Temporal Gyrus showed highly correlated time-courses both in the iERP and the HGA metrics. B-Intra-Parietal Sulcus showed uncorrelated LFP time courses, but correlated HGA time-courses. C-For Pars Opercularis, in the inferior frontal gyrus, the opposite pattern was observed: correlated iERP but uncorrelated HGA. D-In Supramarginal Gyrus, neither metric was correlated across patients.

**Figure 4:**
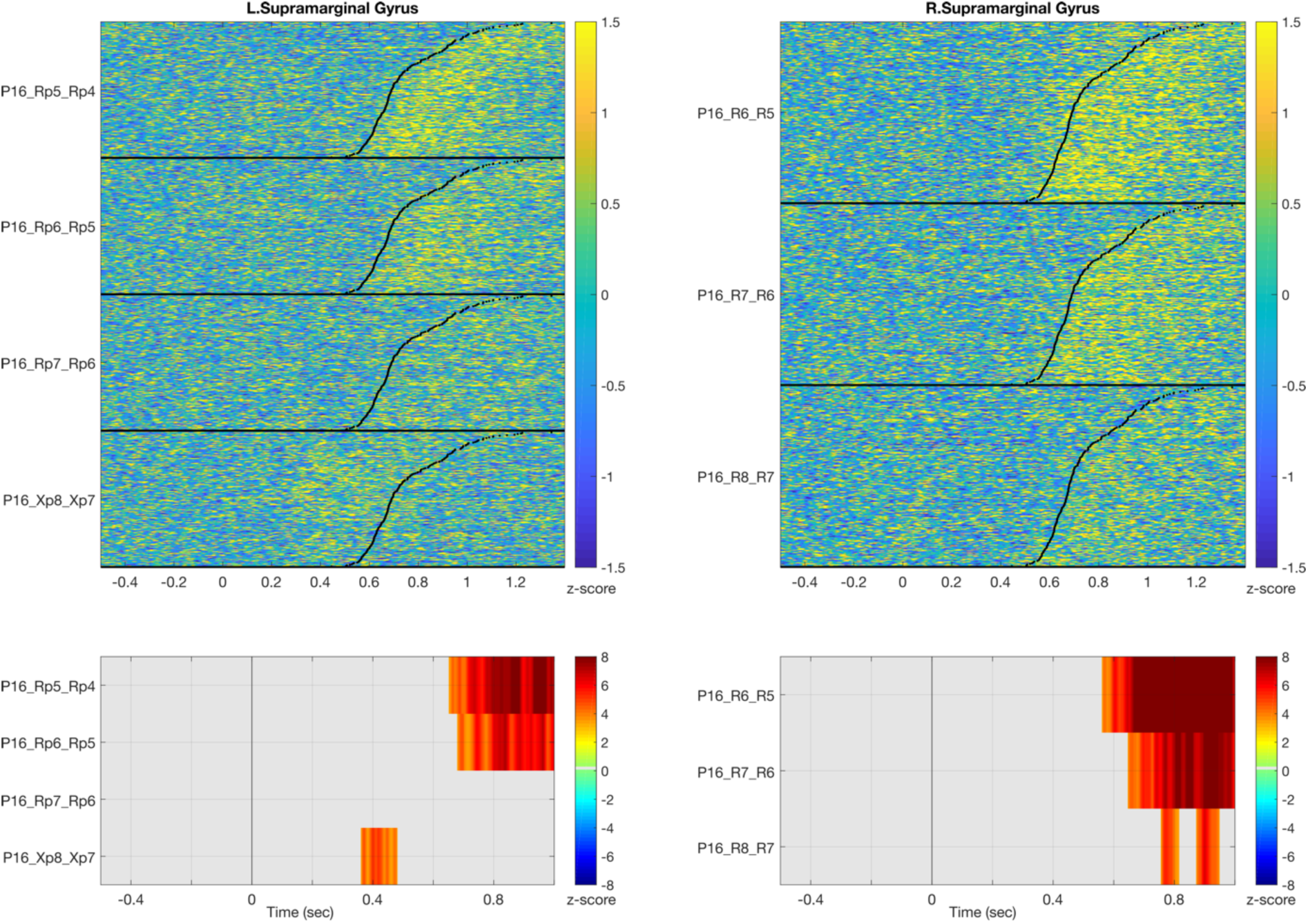
Recordings from the supra sylvian area of SMG in the left (Rp electrode) and right hemisphere (R electrode) and the left retro-sylvian area of SMG (Xp electrode) in Patient 16. Note that the activation post vocalization is recorded bilaterally only in the supra-sylvian area and not in the retro-sylvian area.

The first group included regions for which both iERPs and HGA time courses showed significant correlations across patients and contacts. These “low variability group” include sensory regions such as the superior temporal gyrus (STG) and motor regions such as central sulcus and sub-central gyrus. Figure 3A illustrates the highly concordant activation of STG across contacts and patients, presumably time locked to the patients’ perception of their own voice. That is most clearly seen on the HGA raster-plots (Pt 9 & Pt16). The iERPs components (N600/ P1000) provide additional reliable information on the latency range of involvement of the STG. The temporal pattern of the central sulcus showed a sustained activation long before the onset of articulation (see below 8^th^ panel on Figure 5 and 4^th^ panel on Figure 6).

**Figure 5:**
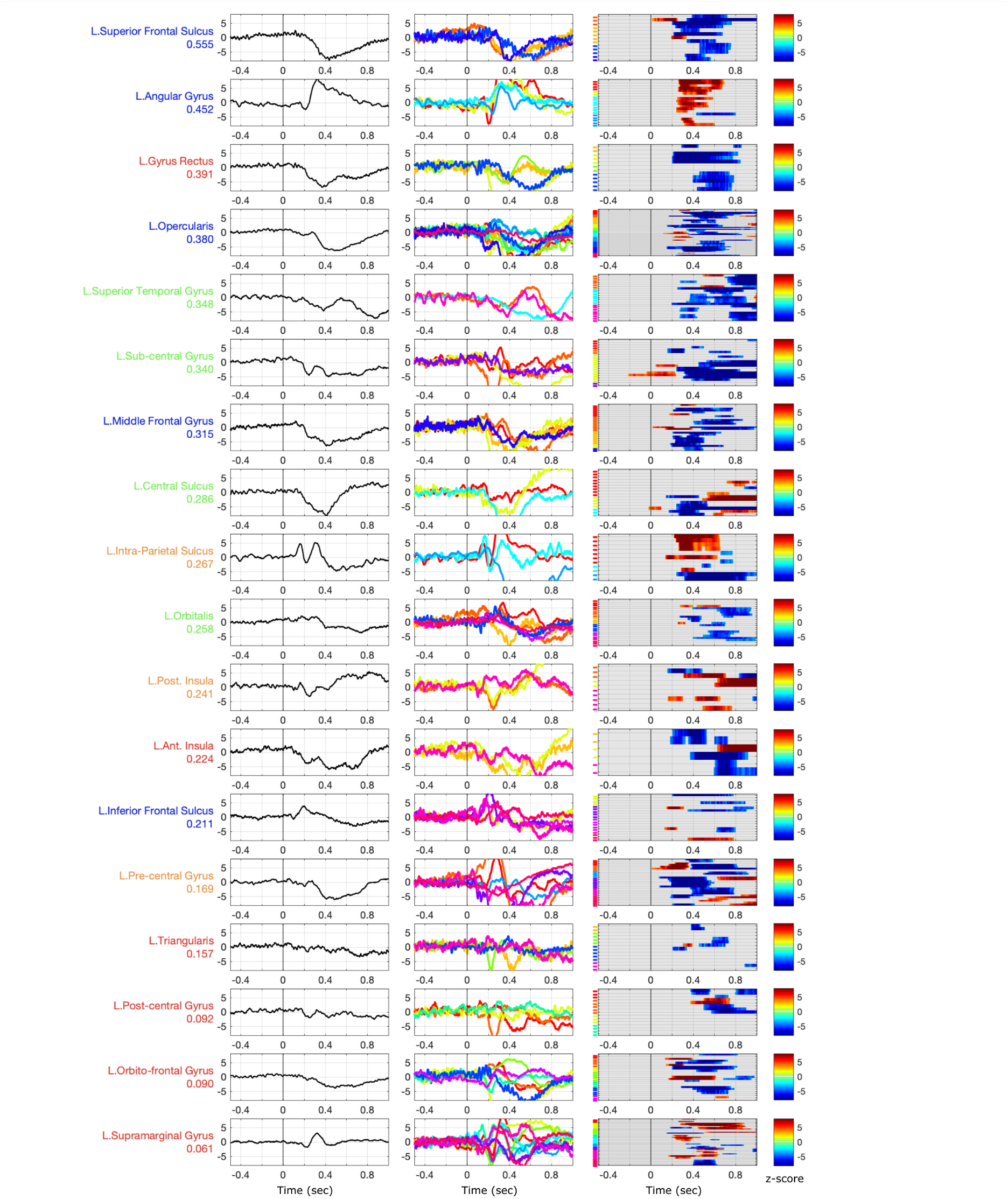
Consistency of iERPs recorded during a word naming tasks across regions of the left dorsal stream. The regions are ordered by decreasing iERP correlation across contacts and patients to highlight the gradient of variability. From left to right columns: grand average, patient averages, and contact-level statistical masks in z-score against baseline.

**Figure 6:**
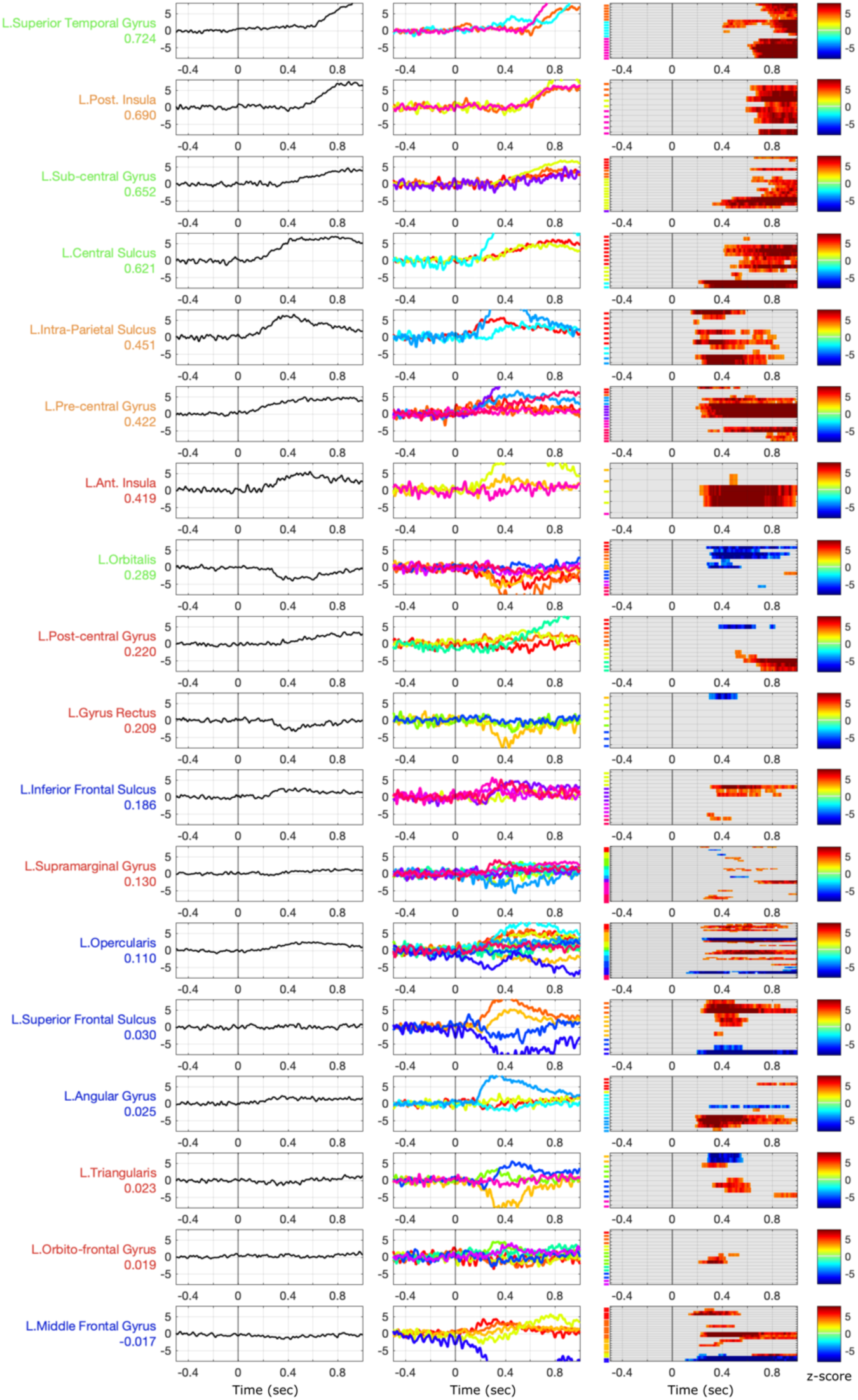
Consistency of High Gamma Activity (HGA) recorded during a word naming tasks across regions of the left dorsal stream. The regions are ordered by decreasing HGA correlation across contacts and patients to highlight the gradient of variability. From left to right columns: grand average, patient averages, and contact-level statistical masks in z-score against baseline.

The second and the third group included regions for which either the HGA or the iERPs time courses, but not both, showed significant correlations across patients and contacts. Consider for example Intra parietal sulcus (IPS; Figure 3B). The average time-course was similar for two of the patients (P01 and P09), an early bi-phasic followed by a more ample response. The third patient (P10) only showed a late ample response, with the opposite polarity. This opposite polarity might reflect different relative positions of the contact probes and the neural generator; this is despite the heuristic procedure we implemented for optimizing similarity across signals irrespective of their sign (see computations described for iERP in *Within-region variability across contacts and patients*). Additionally, the iERP being computed over mono-polar recordings is more sensitive to distant sources that might be present in one patient but not in others. In contrast with the between-patient variability in iERPs, HGA was consistent across patients and even across trials. The single-trial raster plots reveal that IPS activity was linked to stimulus presentation rather than response preparation or execution (Figure 3B right column).

The opposite pattern, consistent iERPs but inconsistent HGAs across patients, was also observed in several regions, for example in Pars Opercularis (Figure 3C). A very systematic iERP response is recorded across the contacts of 11 patients, whereas HGA shows substantial variation across 22 bipolar channels, 18 of which show a significant increase of activity and 6 a significant decrease. HGA has a much smaller power than LFPs, which could contribute to its apparent instability. It is worth noting that the HGA activation starts around 300 ms in all the patients regardless of the response times.

Finally, the fourth group includes areas showing inconsistent activities for both iERP and HGA time-courses. For instance, the supra-marginal gyrus (SMG) presents a range of distinct time-course morphologies and onset times for both neurophysiological measures (Figure 3D). Given the centrality of this region in dorsal stream models of language processing and given that neither of the two metrics show any consistency, a closer examination of the anatomical positions of the contact probes in this region was performed. The post-hoc hypothesis was that the heterogeneity in functional activity may reflect systematic differences in the activity across locations within the SMG region. We distinguished the contacts within SMG according to their anatomical location in supra-sylvian vs. retro-sylvian areas. Interestingly, the contacts located in the supra-sylvian aspect of SMG exhibit a post-vocalization activity, which is also recorded in the homologous right hemisphere region (Figure 4)

Among the contacts located in the retro-sylvian aspect of SMG, iERPs show substantial variability in their temporal courses, yet a component peaking around 300ms is recorded from almost all the contacts. This suggests an early involvement of this region that was not captured by HGA.

The complete list of regions sampled for the study is presented on Figures 5 and 6, sorted according to the variability of their iERP and HGA time-courses, respectively. Angular gyrus (AG) is activated around 250ms reliably across the contacts whereas a prominent slow wave peaking between 400 and 600ms is recorded from the left frontal areas (IFG, IFS, MFG, Pars Opercularis). It is worth noting that the left precentral gyrus does not show a strong consistency across contacts but displays a time course similar to Pars Opercularis’s starting in average around 300ms, long before the average time of overt response (850ms). Interestingly, scattered iERP patterns are observed in 6 out 15 contacts located in the Pars Triangularis.

The analysis of intra-cerebral HGA completes the picture of the naming process across the regions. The motor and auditory regions are consistently activated across their respective contacts. STG and Posterior Insula are involved around the time of the vocalization along with the subcentral gyrus and the central sulcus. The precentral gyrus is involved very early in the process and its activation lasts throughout the vocalization. The timing of HGA fluctuations is quite different within the usual parcellation of Inferior Frontal Gyrus. Pars triangularis displays very short activation or deactivation around 400ms while Pars Opercularis shows a substantial response, with inconsistencies for three out 8 sampled patients. Finally, Pars Orbitalis is mainly deactivated or silent.

In summary, iERP and HGA-consistent patterns of activation are primarily recorded from motor and auditory regions. The timing of the activation seen in these regions varies as function of the patients’ response latencies. The spatio-temporal activity of most of the areas involved in the parieto-frontal language network is consistently captured either by the iERP or the HGA metric. The most notable exceptions are the left Pars Orbitalis, from which no cortical activity is recorded during this task, as well as the left Pars Triangularis and SMG in which there was no reliable activity across patients. Where available, the comparison with response times (RTs) indicates that neural activity is time locked to the stimulus and barely linked to the variation of RTs across trials, except in the supra-sylvian region of SMG.

### Comparison of left and right hemisphere activity in a subsample of the regions

Seven patients were implanted in right hemisphere regions, providing an opportunity for inter-hemispheric comparisons of the variability of functional response in eight regions. The correlation values and their statistical significance are summarized on Figure 7. Three broadly distinct correlation patterns are notable. First, central sulcus showed activity correlated across contacts in both hemispheres and both neurophysiological metrics. Secondly, various regions showed correlated activity in the left hemisphere (for one or both metrics) but not the right hemisphere. This was the case for MFG, Pars Orbitalis, and Pre-central gyrus. Finally, various regions showed correlations that were significant for one or both metrics in the right but not the left hemisphere, in other words there was less variability in the right compared to the left hemisphere. This was the case of Pars triangularis, SMG and Orbito-frontal gyrus.

**Figure 7:**
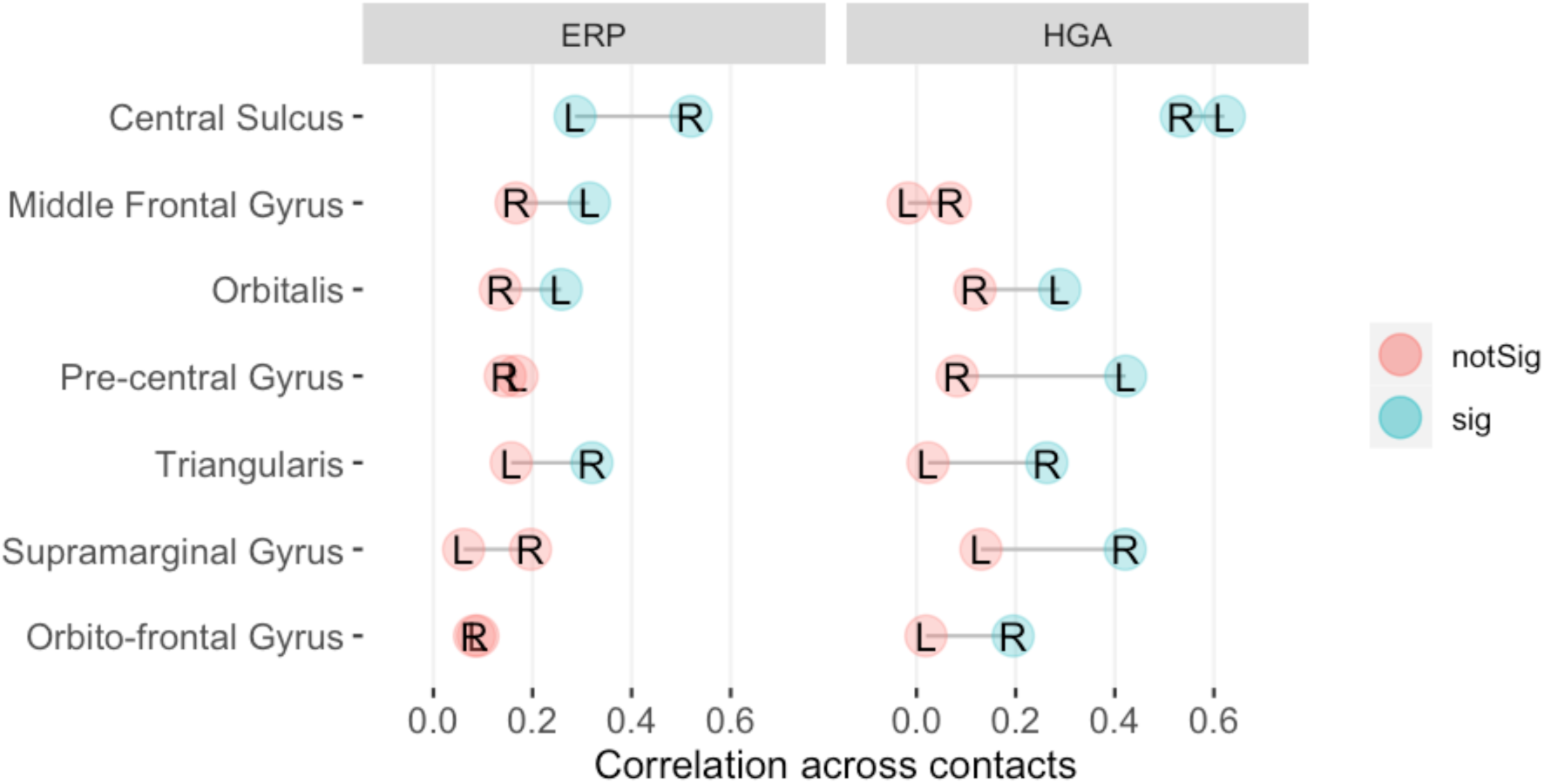
Correlations (values on the x-axis) of neural responses across contacts for regions (y-axis) that were sampled in both hemispheres in the current dataset. L = left; R = right.

## Discussion

We report the recruitment of the different areas composing the dorsal stream during word production. The candidate regions in the temporo-parietal frontal network were defined on anatomical criteria, and the most relevant regions of interest were then selected on the basis of previous functional imaging evidence (e.g. Indefrey, 2011; C. J. Price, 2012). The activity we observed was extensively distributed across the network.

The correlations computed on the iERPs and HGA time-courses revealed contrastive levels of variability. The lowest variability in the patterns of iERP or HGA modulations was observed in auditory cortex (STG) and in some regions of the motor network (central sulcus and subcentral gyrus). These activities were linked to the patients’ responses (auditory feedback and motor articulation). Interestingly, the activity in central sulcus started before and was sustained throughout the overt response, which is consistent with a motor preparation occurring bilaterally. A good spatial agreement between iERP and HGA in STG and the motor cortex is consistent with previous intracranial studies (Edwards et al, 2005; Towle et al, 2008; Sinai et al, 2009; Szurhaj et al, 2006).

Conversely no consistent iERP and HGA deactivation were recorded from the Pars Orbitalis (i.e. in BA 47). This null result extended to the absence or suppression of activity stands in contrast with some studies that showed activation during lexico-semantic control, syntactic processes, and retrieval processes (e.g., Badre et al., 2005; for a review, see Friederici, 2011; Tyler et al., 2011) and the meta-analysis conducted by Binder et al. 2009). This absence of activation can be linked to the simplicity of our task, given item repetitions and the lack of other language processes (e.g., syntactic).

A consistent iERP pattern, implying higher temporal accuracy, is observed in the Angular Gyrus and the Pars Opercularis as well as other frontal regions: MFG, SFG, and SFS. Even if the averaged timing of the engagement of these cortical regions during the naming process must be taken with caution (Dubarry et al., 2017), the semantic processed subserved by the AG activation (Binder et al., 2009; Brownsett & Wise, 2010; A. R. Price et al., 2015) and the word selection subserved by the MFG and SFG (C. J. Price, 2012; S. Riès et al., 2016) overlap with the phonological encoding subserved by the Pars Opercularis (also referred to as Brodmann area BA 44) (for review Friederici et al 2012). The timing of the Pars Opercularis activity before response onset have been previously observed in a large cohort of patients (Nakai et al., 2019) and clearly links its functional role to phonological processing that prepares, rather than executes, the oral response (Flinker et al., 2015).

The variability of the HGA observed in these regions stems from the deactivation or absence of response recorded from some contacts, which could be the result of less active neurons underneath these contacts. Field potential power decays for higher frequencies (1/f) is generally sensitive to the number of active neurons. Another contributing factor is noise. Activities in high gamma frequencies could have poor signal to noise ratio due to contacts being positioned near the skull, which attenuates signal changes (Pesaran et al, 2018). Conversely, a subset of regions involved in speech production were identified only by their significant HGA correlation across patients. This is the case of Precentral Gyrus, whose activity started in average three hundred milliseconds before articulation, confirming that pre-motor areas are involved in an early stage of language production (Fried et al., 1981). In terms of lateralization, (Cogan et al., 2014) argued for a bilateral sub-lexical speech sensory-motor system. Our results show a high reliability in bilateral involvement of central sulcus and left precentral gyrus compared to right Precentral gyrus. These observations were made across patients and the pattern would need to be explored within patient, and thus within trials. With this limitation in mind, the current findings would favour the hypothesis of lateralized sensori-motor transformations (Hickok, 2012; Long et al., 2016).

SMG and Pars Triangularis are two key structures key in the dorsal pathway that belong to the fourth category of regions displaying a high variability across patients in the two metrics. SMG revealed an unexpected heterogeneity of responses, both in LFP and HGA. This disparity can be tentatively linked to the specific location of electrode implantation with a distinction made between its supra- or retro-sylvian parts. HGA time-locked to the articulation is observed only in the supra-sylvian part, bilaterally, whereas scattered activity is recorded from the retro-sylvian region. The activation of SMG in association with speech production and phonological processing has been frequently reported (Fridriksson et al., 2010; Moser et al., 2009). Oberhuber and colleagues (Oberhuber et al., 2016) have proposed a functional parcellation of SMG whereby different types of phonological processing are assigned to four sub-regions, with the anterior ventral subregion being associated with articulatory sequencing. Our data demonstrate that this area is involved in articulatory processing since its activation was time locked with the patient’s response. In addition, the activation of the homotopic contralateral area (in Patient 16) demonstrates that this processing is bilateral. Applying transcranial magnetic stimulation (TMS) Hartwigsen and colleagues (Hartwigsen et al., 2016) showed that a bilateral functional lesion effect induced by TMS entailed a bigger impairment in phonological decision than unilateral stimulations over the left or right SMG.

The findings observed in the Pars Triangularis illustrate well the inconsistency between LFP and HGA, and their variability across the patients. Two patients out of 4 displayed LFP along with increase HGA in one, and deactivation in the others. One patient displayed only significant HGA increase. The activation of Pars Triangularis has been reported during (arguably more demanding) retrieval tasks, such as verbal fluency or verb generation in functional imaging or word-production after morphological inflection (Bourguignon, 2014; Sahin et al., 2009). In functional language mapping, picture naming has often been reported as inducing little HGA activation in the IFG (Arya et al., 2017; Babajani-Feremi et al., 2016; Kunii et al., 2013; Nakai et al., 2017).

## Limitations

Some technical limitations of our analysis must be acknowledged. First, we explored the variability of neural responses both within and across patients. This was done on a medium size group (N=16) although not all patients were recorded in all areas. Future studies based on larger group sizes and focusing on smaller groups of regions of interest should confirm or challenge our conclusions regarding the consistency of activities. As noted in the Methods, the significance of iERP correlations was dependent of a thresholding sign-flipping procedure, on account of potential differences in the relative positions of the generator and the recording contact. The liberal threshold we adopted might have under-estimated the real extent of within– region variability. Secondly, the temporal variability of our analysis stopped at single contacts per region and did not encompass single trials (although single trials were used to compute the time-frequency decompositions). Our interpretations are thus based on averages per contact, which can blur important features of the timing of neural activities (Dubarry et al., 2017). Given the broad spatial coverage reported here (left and part of right dorsal stream), the analysis at the level single trial was left for future, more focused research. Finally, due to clinical constraints during testing, we did not collect response times for every patient; for this reason, we restricted our analysis to stimulus-locked activity.

## Conclusions

Most of the cortical structures known to be involved in word production were identified in this study, either by iERP or HGA. The evidence we report clearly revealed the simultaneous activation of parietal and frontal regions (AG, MFG) which have been linked to semantic processing and word retrieval. Lagging by around 100 ms on the average time-courses was the activation of regions more specialized in phonological processing and articulatory preparation, such as Pars Opercularis, and the Precentral gyrus. These overall timing properties are suggestive of a temporal overlap between regions dedicated to the different sub-processes of word production; they are in line with models that posit interactive processing and the concurrent recruitment of brain areas. Nevertheless, the high variability observed across patients in either one or both neurophysiological metrics confirms that they reflect two facets of the cortical activation subserved by distinct neural mechanisms. Gamma oscillations reflect a competition between excitation and inhibition in local cortical areas while iERP represent the post synaptic activity resulting from the firing of nearby neurons along with remote neurons with afferent inputs into a region (Nunez & Srinivasan, 2006). HGA is more sensitive to changes in the firing rate of the underlying neuronal population than iERP. Its absence does not necessarily mean that there is no synchronized activity, but rather that the threshold of temporally structured spiking is too low to be captured. Our analysis revealed substantial diversity across patients within many of the dorsal stream regions of interest. Figure 8 summarizes the classification into four types of regions, according to the variability observed in the cortical activity. There were sharp distinctions within the left Inferior Frontal Gyrus, with Pars Triangularis and Pars Orbitalis not being systematically recruited during the task. Parietal regions revealed more variability (for either iERP or HGA metrics) than expected based on current cognitive neuroscience models of word production.

**Figure 8:**
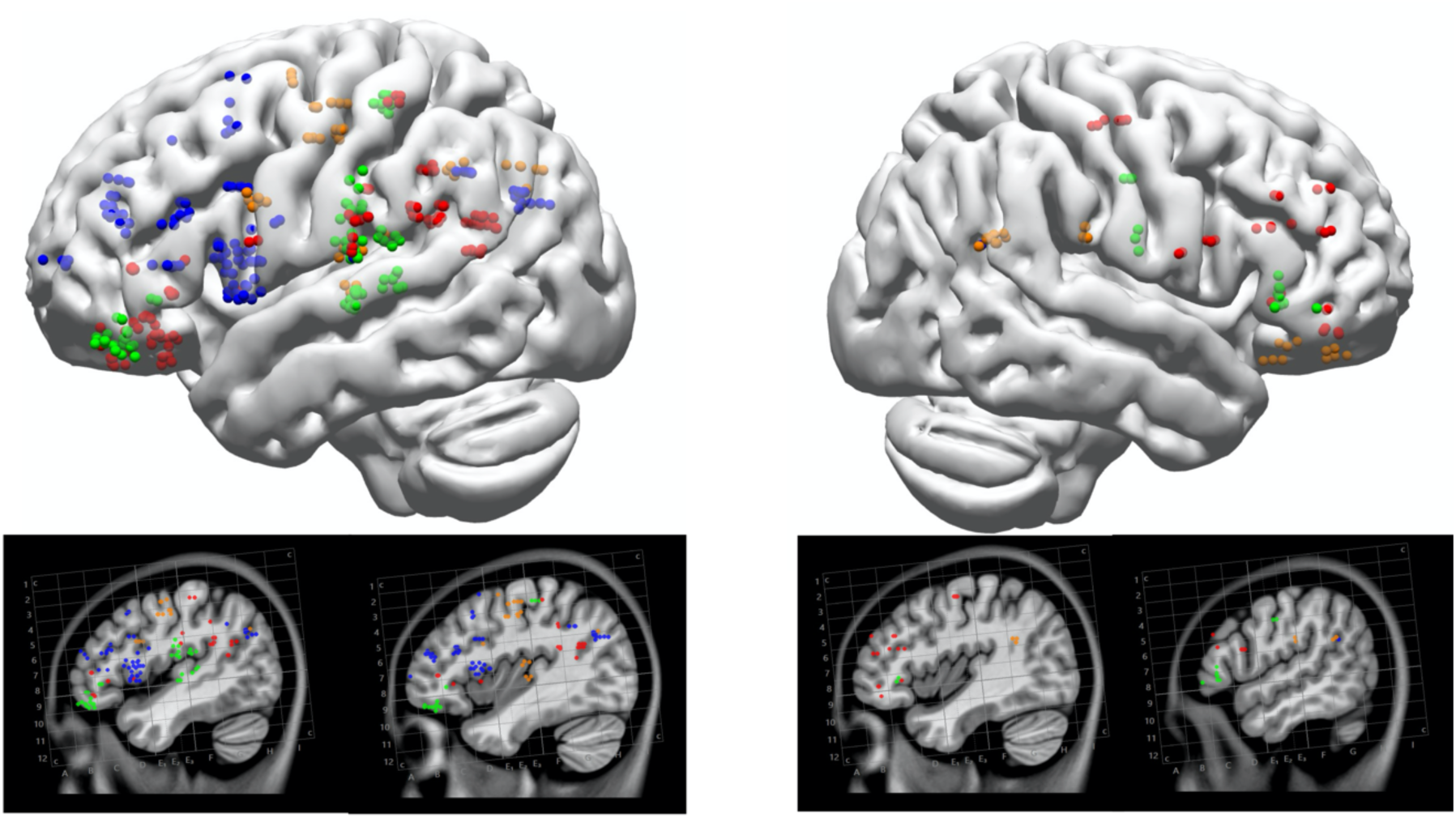
Distribution of the depth electrode coverage across patients in the fronto-parietal dorsal networks. The color of each dot indicates whether the corresponding cortical region showed significantly consistent activities in iERPs and HGA (green), in HGA only (orange), in iERPs only (blue), or in neither of these measures (red).

Tentative interpretations were provided for some of these variations, based on the fine-grained spatial location of the recording contacts within each region. They call for sub-second and sub-region focused explorations of individualized language maps, and for multimodal and individualized language assessments prior to brain surgery involving language cortical areas. Such individualized clinical assessments would benefit from complementing picture naming with alternative experimental tasks designed to maximize the likelihood of activity in the targeted regions.

From such clinical standpoint, the results pose a challenge for language localization at the individual level using picture naming task currently used and indicate the need for specific paradigms of language testing when evaluating language function in clinical settings. This highlights the importance of individualized interpretations of language mapping results, especially in the essential task of word production. While our report does not resolve the issue of the inter-individual variability of SEEG signals, it provides a grounding for future developments in neural signal assessment, clinical interpretations, and neuroscientific theories.

## Data Accessibility statement

The data that support the findings of this study are available on request from the corresponding author. The data are not publicly available due to privacy or ethical restrictions.

## Acknowledgements

We thank all patients for their participation; the Staff of the Cleveland Clinic Epilepsy Center; I Najm, director of the Epilepsy Center for his support; W.Wang & S.Alomar for their help to anatomical data reconstruction and Jean Francois Demonet for his fruitful comments.

## References

1. Anders, R., Llorens, A., Dubarry, A.-S., Trébuchon, A., Liégeois-Chauvel, C., & Alario, F.-X. (2019). Cortical Dynamics of Semantic Priming and Interference during Word Production: An Intracerebral Study. Journal of Cognitive Neuroscience, 1–24. https://doi.org/10.1162/jocn_a_01406

2. Arya, R., Horn, P. S., & Crone, N. E. (2018). ECoG high-gamma modulation versus electrical stimulation for presurgical language mapping. Epilepsy & Behavior, 79, 26–33. https://doi.org/10.1016/j.yebeh.2017.10.044

3. Arya, R., Wilson, J. A., Fujiwara, H., Rozhkov, L., Leach, J. L., Byars, A. W., Greiner, H. M., Vannest, J., Buroker, J., Milsap, G., Ervin, B., Minai, A., Horn, P. S., Holland, K. D., Mangano, F. T., Crone, N. E., & Rose, D. F. (2017). Presurgical language localization with visual naming associated ECoG high-gamma modulation in pediatric drug-resistant epilepsy. Epilepsia, 663–673. https://doi.org/10.1111/epi.13708

4. Babajani-Feremi, A., Narayana, S., Rezaie, R., Choudhri, A. F., Fulton, S. P., Boop, F. A., Wheless, J. W., & Papanicolaou, A. C. (2016). Language mapping using high gamma electrocorticography, fMRI, and TMS versus electrocortical stimulation. Clinical Neurophysiology, 127(3), 1822–1836. https://doi.org/10.1016/j.clinph.2015.11.017

5. Badre, D., Poldrack, R. A., Paré-Blagoev, E. J., Insler, R. Z., & Wagner, A. D. (2005). Dissociable Controlled Retrieval and Generalized Selection Mechanisms in Ventrolateral Prefrontal Cortex. Neuron, 47(6), 907–918. https://doi.org/10.1016/J.NEURON.2005.07.023

6. Bancaud, J., Angelergues, R., Bernouilli, C., Bonis, A., Bordas-Ferrer, M., Bresson, M., Buser, P., Covello, L., Morel, P., Szikla, G., Takeda, A., & Talairach, J. (1969). [Functional stereotaxic exploration (stereo-electroencephalography) in epilepsies]. Revue Neurologique, 120(6), 448.

7. Bédard, C., & Destexhe, A. (2009). Macroscopic Models of Local Field Potentials and the Apparent 1/f Noise in Brain Activity. Biophysical Journal, 96(7), 2589–2603. https://doi.org/10.1016/j.bpj.2008.12.3951

8. Benjamin, C. F. A., Gkiatis, K., Matsopoulos, G. K., & Garganis, K. (2020). Presurgical Language fMRI in Epilepsy: An Introduction. In G. P. D. Argyropoulos (Ed.), Translational Neuroscience of Speech and Language Disorders (pp. 205–239). Springer International Publishing. https://doi.org/10.1007/978-3-030-35687-3_10

9. Berl, M. M., Zimmaro, L. A., Khan, O. I., Dustin, I., Ritzl, E., Duke, E. S., Sepeta, L. N., Sato, S., Theodore, W. H., & Gaillard, W. D. (2014). Characterization of atypical language activation patterns in focal epilepsy. Annals of Neurology, 75(1), 33– 42. https://doi.org/10.1002/ana.24015

10. Binder, J. R., Desai, R. H., Graves, W. W., & Conant, L. L. (2009). Where is the semantic system? A critical review and meta-analysis of 120 functional neuroimaging studies. Cerebral Cortex, 19(12), 2767–2796. https://doi.org/10.1093/cercor/bhp055

11. Bourguignon, N. J. (2014). A rostro-caudal axis for language in the frontal lobe: The role of executive control in speech production. Neuroscience and Biobehavioral Reviews, 47, 431–444. https://doi.org/10.1016/j.neubiorev.2014.09.008

12. Brownsett, S. L. E., & Wise, R. J. S. (2010). The Contribution of the Parietal Lobes to Speaking and Writing. Cerebral Cortex, 20(3), 517–523. https://doi.org/10.1093/cercor/bhp120

13. Buzsáki, G., Anastassiou, C. A., & Koch, C. (2012). The origin of extracellular fields and currents—EEG, ECoG, LFP and spikes. Nature Reviews Neuroscience, 13(6), 407–420. https://doi.org/10.1038/nrn3241

14. Catani, M., & Mesulam, M. (2008). The arcuate fasciculus and the disconnection theme in language and aphasia: History and current state. Cortex, 44(8), 953–961. https://doi.org/10.1016/j.cortex.2008.04.002

15. Caune, V., Ranta, R., Le Cam, S., Hofmanis, J., Maillard, L., Koessler, L., & Louis-Dorr, V. (2014). Evaluating dipolar source localization feasibility from intracerebral SEEG recordings. NeuroImage, 98, 118–133. https://doi.org/10.1016/J.NEUROIMAGE.2014.04.058

16. Chauvel, P., Gonzalez-Martinez, J., & Bulacio, J. (2019). Presurgical intracranial investigations in epilepsy surgery. Handbook of Neurology. Elsevier.

17. Cogan, G. B., Thesen, T., Carlson, C., Doyle, W., Devinsky, O., & Pesaran, B. (2014). Sensory–motor transformations for speech occur bilaterally. Nature, 507(7490), 94–98. https://doi.org/10.1038/nature12935

18. Collins, D. L., Zijdenbos, A. P., Kollokian, V., Sled, J. G., Kabani, N. J., Holmes, C. J., & Evans, A. C. (1998). Design and construction of a realistic digital brain phantom. IEEE Transactions on Medical Imaging, 17(3), 463–468. https://doi.org/10.1109/42.712135

19. D’Esposito, M. (2019). Are individual differences in human brain organization measured with functional MRI meaningful? Proceedings of the National Academy of Sciences, 116(45), 22432–22434. https://doi.org/10.1073/pnas.1915982116

20. Dubarry, A.-S., Llorens, A., Trébuchon, A., Carron, R., Liégeois-Chauvel, C., Bénar, C. G., & Alario, F. X. (2017). Estimating Parallel Processing in a Language Task Using Single-Trial Intracerebral Electroencephalography. Psychological Science, 28(4), 414–426. https://doi.org/10.1177/0956797616681296

21. Edwards, E., Soltani, M., Kim, W., Dalal, S. S., Nagarajan, S. S., Berger, M. S., & Knight, R. T. (2009). Comparison of Time-Frequency Responses and the Event-Related Potential to Auditory Speech Stimuli in Human Cortex. Journal of Neurophysiology, 102(1), 377–386. https://doi.org/10.1152/jn.90954.2008

22. Fedorenko, E., Behr, M. K., & Kanwisher, N. (2011). Functional specificity for high-level linguistic processing in the human brain. Proceedings of the National Academy of Sciences, 108(39), 16428–16433. https://doi.org/10.1073/pnas.1112937108

23. Flinker, A., Korzeniewska, A., Shestyuk, A. Y., Franaszczuk, P. J., Dronkers, N. F., Knight, R. T., & Crone, N. E. (2015). Redefining the role of Broca’s area in speech. Proceedings of the National Academy of Sciences, 112(9), 2871–2875. https://doi.org/10.1073/pnas.1414491112

24. Flinker, A., Piai, V., & Knight, R. (2018). Intracranial electrophysiology in language research. The Oxford Handbook of Psycholinguistics (2 Ed.). https://doi.org/10.1093/oxfordhb/9780198568971.001.0001

25. Fridriksson, J., Kjartansson, O., Morgan, P. S., Hjaltason, H., Magnusdottir, S., Bonilha, L., & Rorden, C. (2010). Impaired Speech Repetition and Left Parietal Lobe Damage. Journal of Neuroscience, 30(33), 11057–11061. https://doi.org/10.1523/JNEUROSCI.1120-10.2010

26. Fried, I., Ojemann, G. A., & Fetz, E. E. (1981). Language-Related Potentials Specific to Human Language Cortex. Science, 212(4492), 353–356. https://doi.org/10.1126/science.7209537

27. Friederici, A. D. (2011). The brain basis of language processing: From structure to function. Physiological Reviews, 91(4), 1357–1392. https://doi.org/10.1152/physrev.00006.2011

28. Gleichgerrcht, E., Fridriksson, J., & Bonilha, L. (2015). Neuroanatomical foundations of naming impairments across different neurologic conditions. Neurology, 85(3), 284–292. https://doi.org/10.1212/WNL.0000000000001765

29. González-Martínez, J. (2015). Convergence of stereotactic surgery and epilepsy: The stereoelectroencephalography method. Neurosurgery, 62(CN_suppl_1), 117–122. https://doi.org/10.1227/NEU.0000000000000787

30. Gonzalez-Martinez, J., Mullin, J., Vadera, S., Bulacio, J., Hughes, G., Jones, S., Enatsu, R., & Najm, I. (2014). Stereotactic placement of depth electrodes in medically intractable epilepsy. Journal of Neurosurgery, 120(3), 639–644. https://doi.org/10.3171/2013.11.jns13635

31. Gordon, E. M., Laumann, T. O., Gilmore, A. W., Newbold, D. J., Greene, D. J., Berg, J. J., Ortega, M., Hoyt-Drazen, C., Gratton, C., Sun, H., Hampton, J. M., Coalson, R. S., Nguyen, A. L., McDermott, K. B., Shimony, J. S., Snyder, A. Z., Schlaggar, B. L., Petersen, S. E., Nelson, S. M., & Dosenbach, N. U. F. (2017). Precision Functional Mapping of Individual Human Brains. Neuron, 95(4), 791–807.e7. https://doi.org/10.1016/j.neuron.2017.07.011

32. Hartwigsen, G., Weigel, A., Schuschan, P., Siebner, H. R., Weise, D., Classen, J., & Saur, D. (2016). Dissociating Parieto-Frontal Networks for Phonological and Semantic Word Decisions: A Condition-and-Perturb TMS Study. Cerebral Cortex, 26(6), 2590–2601. https://doi.org/10.1093/cercor/bhv092

33. Hickok, G. (2012). Computational neuroanatomy of speech production. Nature Reviews Neuroscience, 13(2), 135–145. https://doi.org/10.1038/nrn3158

34. Indefrey, P. (2011). The Spatial and Temporal Signatures of Word Production Components: A Critical Update. Frontiers in Psychology, 2(1–2), 255. https://doi.org/10.3389/fpsyg.2011.00255

35. Johnson, M. A., Thompson, S., Gonzalez-Martinez, J., Park, H.-J., Bulacio, J., Najm, I., Kahn, K., Kerr, M., Sarma, S. V., & Gale, J. T. (2014). Performing Behavioral Tasks in Subjects with Intracranial Electrodes. Journal of Visualized Experiments, 92, e51947. https://doi.org/10.3791/51947

36. Kunii, N., Kamada, K., Ota, T., Greenblatt, R. E., Kawai, K., & Saito, N. (2013). The dynamics of language-related high-gamma activity assessed on a spatially-normalized brain. Clinical Neurophysiology, 124(1), 91–100. https://doi.org/10.1016/j.clinph.2012.06.006

37. Lachaux, J. P., Axmacher, N., Mormann, F., Halgren, E., & Crone, N. E. (2012). High-frequency neural activity and human cognition: Past, present and possible future of intracranial EEG research. Progress in Neurobiology, 98(3), 279–301. https://doi.org/10.1016/j.pneurobio.2012.06.008

38. Levelt, W. J. M., Roelofs, A., & Meyer, A. S. (1999). A theory of lexical access in speech production. Behavioral and Brain Sciences, 22(1), 1–75. https://doi.org/10.1017/S0140525X99001776

39. Llorens, A., Dubarry, A.-S., Trébuchon Da Fonseca, A., Chauvel, P., Alario, F.-X., & Liégeois-Chauvel, C. (2016). Contextual modulation of hippocampal activity during picture naming. Brain and Language, 159, 92–101. https://doi.org/10.1016/j.bandl.2016.05.011

40. Llorens, A., Trébuchon, A., Liégeois-Chauvel, C., & Alario, F.-X. (2011). Intra-cranial recordings of brain activity during language production. Frontiers in Psychology, 2(DEC), 1–12. https://doi.org/10.3389/fpsyg.2011.00375

41. Logothetis, N. K., Pauls, J., Augath, M., Trinath, T., & Oeltermann, A. (2001). Neurophysiological investigation of the basis of the fMRI signal. Nature, 412(6843), 150–157. https://doi.org/10.1038/35084005

42. Long, M. A., Katlowitz, K. A., Svirsky, M. A., Clary, R. C., Byun, T. M. A., Majaj, N., Oya, H., Howard, M. A., & Greenlee, J. D. W. (2016). Functional Segregation of Cortical Regions Underlying Speech Timing and Articulation. Neuron, 89(6), 1187–1193. https://doi.org/10.1016/j.neuron.2016.01.032

43. Lopes da Silva, F. (1991). Neural mechanisms underlying brain waves: From neural membranes to networks. Electroencephalography and Clinical Neurophysiology, 79(2), 81–93. https://doi.org/10.1016/0013-4694(91)90044-5

44. Mahowald, K., & Fedorenko, E. (2016). Reliable individual-level neural markers of high-level language processing: A necessary precursor for relating neural variability to behavioral and genetic variability. NeuroImage, 139, 74–93. https://doi.org/10.1016/j.neuroimage.2016.05.073

45. McKinnon, E. T., Fridriksson, J., Basilakos, A., Hickok, G., Hillis, A. E., Spampinato, M. V., Gleichgerrcht, E., Rorden, C., Jensen, J. H., Helpern, J. A., & Bonilha, L. (2018). Types of naming errors in chronic post-stroke aphasia are dissociated by dual stream axonal loss. Scientific Reports, 8(1), 14352. https://doi.org/10.1038/s41598-018-32457-4

46. Miller, M. B., Donovan, C.-L., Van Horn, J. D., German, E., Sokol-Hessner, P., & Wolford, G. L. (2009). Unique and persistent individual patterns of brain activity across different memory retrieval tasks. NeuroImage, 48(3), 625–635. https://doi.org/10.1016/j.neuroimage.2009.06.033

47. Moser, D., Baker, J. M., Sanchez, C. E., Rorden, C., & Fridriksson, J. (2009). Temporal Order Processing of Syllables in the Left Parietal Lobe. Journal of Neuroscience, 29(40), 12568–12573. https://doi.org/10.1523/JNEUROSCI.5934-08.2009

48. Mukamel, R., & Fried, I. (2012). Human Intracranial Recordings and Cognitive Neuroscience. Annual Review of Psychology, 63(1), 511–537. https://doi.org/10.1146/annurev-psych-120709-145401

49. Munding, D., Dubarry, A. S., & Alario, F.-X. (2016). On the cortical dynamics of word production: A review of the MEG evidence. Language, Cognition and Neuroscience, 31(4), 441–462. https://doi.org/10.1080/23273798.2015.1071857

50. Nakai, Y., Jeong, J. W., Brown, E. C., Rothermel, R., Kojima, K., Kambara, T., Shah, A., Mittal, S., Sood, S., & Asano, E. (2017). Three- and four-dimensional mapping of speech and language in patients with epilepsy. Brain, 140(5), 1351– 1370. https://doi.org/10.1093/brain/awx051

51. Nakai, Y., Sugiura, A., Brown, E. C., Sonoda, M., Jeong, J., Rothermel, R., Luat, A. F., Sood, S., & Asano, E. (2019). Four-dimensional functional cortical maps of visual and auditory language: Intracranial recording. Epilepsia, 60(2), 255–267. https://doi.org/10.1111/epi.14648

52. Nourski, K. V., Steinschneider, M., Rhone, A. E., Oya, H., Kawasaki, H., Howard, M. A., & McMurray, B. (2015). Sound identification in human auditory cortex: Differential contribution of local field potentials and high gamma power as revealed by direct intracranial recordings. Brain and Language, 148, 37–50. https://doi.org/10.1016/j.bandl.2015.03.003

53. Nunez, P. L., & Srinivasan, Ramesh. (2006). Electric fields of the brain: The neurophysics of EEG. Oxford University Press.

54. Oberhuber, M., Hope, T. M. H., Seghier, M. L., Parker Jones, O., Prejawa, S., Green, D. W., & Price, C. J. (2016). Four Functionally Distinct Regions in the Left Supramarginal Gyrus Support Word Processing. Cerebral Cortex, 26(11), 4212–4226. https://doi.org/10.1093/cercor/bhw251

55. Parlatini, V., Radua, J., Dell’Acqua, F., Leslie, A., Simmons, A., Murphy, D. G., Catani, M., & Thiebaut de Schotten, M. (2017). Functional segregation and integration within fronto-parietal networks. NeuroImage, 146, 367–375. https://doi.org/10.1016/j.neuroimage.2016.08.031

56. Parvizi, J., & Kastner, S. (2018). Promises and limitations of human intracranial electroencephalography. Nature Neuroscience, 21(4), 474–483. https://doi.org/10.1038/s41593-018-0108-2

57. Petrides, M. (2014). Neuroanatomy of Language Regions of the Human Brain (First edition). Elsevier/AP, Academic Press is an imprint of Elsevier.

58. Piai, V., Anderson, K. L., Lin, J. J., Dewar, C., Parvizi, J., Dronkers, N. F., & Knight, R. T. (2016). Direct brain recordings reveal hippocampal rhythm underpinnings of language processing. Proceedings of the National Academy of Sciences. https://doi.org/10.1073/pnas.1603312113

59. Price, A. R., Bonner, M. F., Peelle, J. E., & Grossman, M. (2015). Converging evidence for the neuroanatomic basis of combinatorial semantics in the angular gyrus. The Journal of Neuroscience : The Official Journal of the Society for Neuroscience, 35(7), 3276–3284. https://doi.org/10.1523/JNEUROSCI.3446-14.2015

60. Price, C. J. (2012). A review and synthesis of the first 20 years of PET and fMRI studies of heard speech, spoken language and reading. NeuroImage, 62(2), 816–847. https://doi.org/10.1016/j.neuroimage.2012.04.062

61. Rapp, B., & Goldrick, M. (2000). Discreteness and interactivity in spoken word production. Psychological Review, 107(3), 460–499. https://doi.org/10.1037/0033-295X.107.3.460

62. Ray, S., & Maunsell, J. H. R. (2011). Different Origins of Gamma Rhythm and High-Gamma Activity in Macaque Visual Cortex. PLoS Biology, 9(4), e1000610. https://doi.org/10.1371/journal.pbio.1000610

63. Riès, S., Dronkers, N. F., & Knight, R. T. (2016). Choosing words: Left hemisphere, right hemisphere, or both? Perspective on the lateralization of word retrieval. Annals of the New York Academy of Sciences, 1369(1), 111–131. https://doi.org/10.1111/nyas.12993

64. Riès, S. K., Dhillon, R. K., Clarke, A., King-Stephens, D., Laxer, K. D., Weber, P. B., Kuperman, R. A., Auguste, K. I., Brunner, P., Schalk, G., Lin, J. J., Parvizi, J., Crone, N. E., Dronkers, N. F., & Knight, R. T. (2017). Spatiotemporal dynamics of word retrieval in speech production revealed by cortical high-frequency band activity. Proceedings of the National Academy of Sciences, 114(23), E4530–E4538. https://doi.org/10.1073/pnas.1620669114

65. Roelofs, A. (2014). A dorsal-pathway account of aphasic language production: The WEAVER++/ARC model. Cortex, 59, 33–48. https://doi.org/10.1016/j.cortex.2014.07.001

66. Sahin, N. T., Pinker, S., Cash, S. S., Schomer, D., & Halgren, E. (2009). Sequential processing of lexical, grammatical, and phonological information within broca’s area. Science, 326(5951), 445–449. https://doi.org/10.1126/science.1174481

67. Sanfratello, L., Caprihan, A., Stephen, J. M., Knoefel, J. E., Adair, J. C., Qualls, C., Lundy, S. L., & Aine, C. J. (2014). Same task, different strategies: How brain networks can be influenced by memory strategy. Human Brain Mapping, 35(10), 5127–5140. https://doi.org/10.1002/hbm.22538

68. Saur, D., Kreher, B. W., Schnell, S., Kummerer, D., Kellmeyer, P., Vry, M.-S., Umarova, R., Musso, M., Glauche, V., Abel, S., Huber, W., Rijntjes, M., Hennig, J., & Weiller, C. (2008). Ventral and dorsal pathways for language. Proceedings of the National Academy of Sciences, 105(46), 18035–18040. https://doi.org/10.1073/pnas.0805234105

69. Schwartz, M. F., Faseyitan, O., Kim, J., & Coslett, H. B. (2012). The dorsal stream contribution to phonological retrieval in object naming. Brain, 135(12), 3799– 3814. https://doi.org/10.1093/brain/aws300

70. Seghier, M. L., Lee, H. L., Schofield, T., Ellis, C. L., & Price, C. J. (2008). Inter-subject variability in the use of two different neuronal networks for reading aloud familiar words. NeuroImage, 42(3), 1226–1236. https://doi.org/10.1016/j.neuroimage.2008.05.029

71. Seghier, M. L., & Price, C. J. (2018). Interpreting and Utilising Intersubject Variability in Brain Function. Trends in Cognitive Sciences, 22(6), 517–530. https://doi.org/10.1016/j.tics.2018.03.003

72. Seitzman, B. A., Gratton, C., Laumann, T. O., Gordon, E. M., Adeyemo, B., Dworetsky, A., Kraus, B. T., Gilmore, A. W., Berg, J. J., Ortega, M., Nguyen, A., Greene, D. J., McDermott, K. B., Nelson, S. M., Lessov-Schlaggar, C. N., Schlaggar, B. L., Dosenbach, N. U. F., & Petersen, S. E. (2019). Trait-like variants in human functional brain networks. Proceedings of the National Academy of Sciences, 116(45), 22851–22861. https://doi.org/10.1073/pnas.1902932116

73. Snodgrass, J. G., & Vanderwart, M. (1980). A standardized set of 260 pictures: Norms for name agreement, image agreement, familiarity, and visual complexity. Journal of Experimental Psychology. Human Learning and Memory, 6(2), 174–215.

74. Strijkers, K., & Costa, A. (2016). The cortical dynamics of speaking: Present shortcomings and future avenues. Language, Cognition and Neuroscience, 31(4), 484–503. https://doi.org/10.1080/23273798.2015.1120878

75. Tadel, F., Baillet, S., Mosher, J. C., Pantazis, D., & Leahy, R. M. (2011). Brainstorm: A User-Friendly Application for MEG/EEG Analysis. Computational Intelligence and Neuroscience, 2011, 1–13. https://doi.org/10.1155/2011/879716

76. Talairach, J., & Tournoux, P. (1988). Co-Planar Stereotaxic Atlases of the Human Brain: 3-Dimensional Proportional System – An Approach to Cerebral Imaging. Thieme Medical Publishers.

77. Tallon-Baudry, C., & Bertrand, O. (1999). Oscillatory gamma activity in humans and its role in object representation. Trends in Cognitive Sciences, 3(4), 151–162. https://doi.org/10.1016/S1364-6613(99)01299-1

78. Trebuchon, A., Liégeois-Chauvel, C., Gonzalez-Martinez, J. A., & F.-Xavier Alario. (2020). Contributions of electrophysiology for identifying cortical language systems in patients with epilepsy. Epilepsy & Behavior, 112, 107407. https://doi.org/10.1016/j.yebeh.2020.107407

79. Tyler, L. K., Marslen-Wilson, W. D., Randall, B., Wright, P., Devereux, B. J., Zhuang, J., Papoutsi, M., & Stamatakis, E. A. (2011). Left inferior frontal cortex and syntax: Function, structure and behaviour in patients with left hemisphere damage. Brain, 134(2), 415–431. https://doi.org/10.1093/brain/awq369

80. Tzourio-Mazoyer, N., Perrone-Bertolotti, M., Jobard, G., Mazoyer, B., & Baciu, M. (2017). Multi-factorial modulation of hemispheric specialization and plasticity for language in healthy and pathological conditions: A review. Cortex, 86, 314–339. https://doi.org/10.1016/j.cortex.2016.05.013

81. Ueno, T., Saito, S., Rogers, T. T., & Lambon Ralph, M. A. (2011). Lichtheim 2: Synthesizing aphasia and the neural basis of language in a neurocomputational model of the dual dorsal-ventral language pathways. Neuron, 72(2), 385–396. https://doi.org/10.1016/j.neuron.2011.09.013

82. Warren, J. E., Wise, R. J. S., & Warren, J. D. (2005). Sounds do-able: Auditory–motor transformations and the posterior temporal plane. Trends in Neurosciences, 28(12), 636–643. https://doi.org/10.1016/J.TINS.2005.09.010

